# Differential Effects of RASA3 Mutations on Hematopoiesis are Profoundly Influenced by Genetic Background and Molecular Variant

**DOI:** 10.1101/2020.05.14.095745

**Authors:** Raymond F. Robledo, Steven L. Ciciotte, Joel H. Graber, Yue Zhao, Amy J. Lambert, Babette Gwynn, Nathaniel J. Maki, Lionel Blanc, Luanne L. Peters

## Abstract

Studies of the severely pancytopenic *scat* mouse model first demonstrated the crucial role of RASA3, a dual RAS and RAP GTPase activating protein (GAP), in hematopoiesis. RASA3 is required for survival *in utero*; germline deletion is lethal at E12.5-13.5 due to severe hemorrhage and decreased fetal liver erythropoiesis. Conditional deletion in hematopoietic stem and progenitor cells (HSPCs) using *Vav*-*Cre* recapitulates the null phenotype demonstrating that RASA3 is required at the stem and progenitor level to maintain blood vessel development and integrity and effective blood production. In adults, bone marrow blood cell production and spleen stress erythropoiesis are suppressed significantly upon induction of RASA3 deficiency, leading to pancytopenia and death within two weeks. Notably, RASA3 missense mutations in mouse models *scat* (G125V) and *hlb381* (H794L) show dramatically different hematopoietic consequences specific to both genetic background and molecular variant. Global transcriptomic studies in *scat* suggest potential targets to ameliorate disease progression.

**Author Summary:** Hematopoiesis is the process by which blood cells are formed. The individual must have a normal complement of red blood cells to prevent anemia, platelets to control bleeding, and white blood cells to maintain immune functions. All blood cells are derived from hematopoietic stem cells that differentiate into progenitor cells that then develop into mature circulating cells. We studied several mouse strains carrying different mutations in RASA3. We show that RASA3 is required at the earliest stages of blood formation, the stem and progenitor cells, and that the complement of genes other than RASA3, or the genetic background of the mutant strain, profoundly alters the overall effect on blood formation. Further, the molecular nature of the mutation in RASA3 also has a profound and independent effect on overall blood formation. One strain, designated *scat*, suffers cyclic anemia characterized by severe anemic crisis episodes interspersed with remissions where the anemia significantly improves. Comparison of *scat* crisis and remission hematopoietic stem and progenitor cells reveals striking differences in gene expression. Analyses of these expression differences provide clues to processes that potentially drive improvement of anemia in *scat* and provide new avenues to pursue in future studies to identify novel therapeutics for anemia.

## Introduction

RASA3 (RAS p21 protein activator 3), also called GAPIII or IP4BP (inositol 1,3,4,5-tetrakisphosphate, IP4, binding protein), is a member of the GAP1 family of RAS-GTPase-activating proteins (GAPs) that also includes RASA2 (GAP1^m^), RASA4 (CAPRI) and RASAL1 (RASAL) [1]. The RAS superfamily of small GTPases includes the subfamily RAP. Small GTPases act as molecular switches, cycling between active GTP-bound and inactive GDP-bound forms. They are activated by guanine nucleotide exchange factors (GEFs), which stimulate GTP loading, and inactivated by GAPs, which accelerate GTP hydrolysis. Membrane localization is required for GAP activity for all GAP1 family members, which are highly conserved and share a common structure consisting of dual N-terminal C2 domains, a central catalytic GAP domain, and C-terminal pleckstrin homology (PH) and Bruton tyrosine kinase (Btk) domains [2–4].

With the exception of RASA2, members of the GAP1 subfamily are dual RAS and RAP GAPs [4–6]. RAS is a critical mediator of cytokine-dependent signaling in erythropoiesis (erythropoietin, Epo; stem cell factor, SCF) [7–9] and thrombopoiesis (thrombopoietin, TPO) [10], transmitting signals to multiple effector pathways that regulate cell proliferation, differentiation, survival, adhesion, and actin cytoskeleton organization. RAP proteins are highly expressed in endothelial cells, leukocytes, and platelets and play major roles in cell polarity, adhesion, and movement [11].

A major role of RASA3 in hematopoiesis was first identified upon positional cloning of the co-isogenic autosomal recessive mouse mutation, *scat* (severe combined anemia and thrombocytopenia) [12]. The *scat* phenotype, in addition to severe anemia and thrombocytopenia, includes significant leukopenia as well. The *scat* disease progresses episodically, with periods of severe crisis interspersed with one or two periods of remission. Notably, full remission as presented in the original description [13] of the *scat* disease, is now exceedingly rare. Instead, partial remission occurs, which is restricted to the erythroid component – leukocyte and platelet numbers do not recover. This change in *scat* disease progression is likely due to genetic drift over the thirty years since its original description, admixture of two BALBc/By sublines (see Methods), and environmental changes (e. g., changes in animal room health status standards established and maintained since the late 1980s). The RASA3 mutation (G125V) in *scat* causes mislocalization of RASA3 to the cytosol, abrogating RASA3 GAP activity and increasing active RAS levels in *scat* erythroid cells. Increased active RAS is associated with delayed terminal erythroid differentiation [8, 9]. Indeed, terminal erythropoiesis in *scat* is significantly delayed at the poly- and orthochromatophilic developmental stages [12].

Additional studies in mouse models have defined RAP-dependent RASA3 functions *in vivo*. Studies in *hlb381*, a recessive chemically (N-Ethyl-N-Nitrosourea, ENU) induced model carrying a different missense mutation (H794L) in *Rasa3*, confirmed RASA3 as a critical inhibitor of RAP1-dependent integrin signaling and platelet activation [14]. Defects in adhesion, as reflected by severe hemorrhage and reduced numbers of adherens junctions between endothelial cells in vessels of the brain, were noted in the first knockout of *Rasa3* in mice where exons 11 and 12 within the central catalytic GAP domain were replaced with a neomycin cassette; the model was embryonic lethal at E12.5-13.5 [15]. Studies in bone marrow transplant and conditional catalytically inactive RASA3 models revealed RAP1-dependent defects in megakaryocyte integrin signaling and structure (disorganized actin cytoskeleton), platelet adhesion and activation, endothelial cell adhesion and vascular lumen integrity [16, 17].

Here, we sought to further analyze the role of RASA3 in hematopoiesis using conditional knockout models to determine the status of HSPCs in RASA3 deficiency, to elucidate mechanistic differences underlying H794L (*hlb381*) and G125V (*scat*), and to perform RNAseq studies to generate hypotheses regarding the progression of the *scat* disease from periods of crisis to partial remission. Our results show that (1) there is a strict requirement for RASA3 throughout development and into adulthood at the level of HSPCs in order for hematopoiesis to progress normally, (2) adult bone marrow hematopoiesis and stress erythropoiesis in the spleen are suppressed upon induction of RASA3 deficiency, (3) genetic background and the nature of the molecular variant profoundly influence the RASA3 deficient phenotype, and (4) global transcriptomics reveals massive expression differences in *scat vs.* wild type (WT) hematopoietic tissues and cells, and generates novel hypotheses on potential mechanisms leading to spontaneous disease amelioration (partial remission).

## Results

### Mice and breeding

Table S1 lists strains of mice used in this study. A schematic of the overall breeding strategy used to generate *Rasa3* alleles is shown in Fig S1. Except where otherwise noted, all studies were performed using mice on a hybrid B6;129 genetic background. Mice designated as controls are Cre-negative fl/− and fl/fl and/or mice carrying a WT allele regardless of Cre genotype. Mutant mice are Cre-positive fl/− or fl/fl.

*Rasa3* null mice are embryonic lethal. We generated a conditional knockout (cKO) allele for *Rasa3* on a hybrid B6;129 genetic background using the *Rasa3*^tm1a(KOMP)Wtsi^ knockout-first, promoter-driven conditional ready construct generated by the Knock-Out Mouse Project (KOMP) in which exon 3, required for RASA3 GAP activity, is flanked by *loxP* sites (Fig 1A). Correctly targeted ES cells were identified by Southern blotting of genomic DNA using 5- and 3-prime flanking probes (Fig 1B) generated by PCR (primer sequences and expected product sizes provided in Table S2). Heterozygous *Rasa3*^*cKO*/+^ offspring of male chimeras and C57BL/6J (B6J) females were identified by PCR genotyping (Fig 1C) and mated to *ACT-FLPe* transgenic mice to remove the promoter-driven cassette to produce *Rasa3*^*fl*/+^ mice. *Rasa3*^*fl*/+^ mice were bred to *CMV-Cre* transgenic mice to produce the germline *Rasa3* null allele and subsequently backcrossed to B6J mice to remove *CMV-Cre* (Fig 1D). The resultant *Rasa3*^+/−^ mice appeared normal in all respects and were intercrossed. No *Rasa3*^−/−^ pups were obtained at birth, while 16 *Rasa3*^+/−^ (57%) and 12 *Rasa3*^+/+^ (43%) progeny were obtained, indicating embryonic lethality in the absence of RASA3. Studies performed by the International Mouse Phenotyping Consortium (IMPC) confirmed embryonic lethality at E12.5-E13.5 in *Rasa3*^−/−^ null mice on the inbred C57BL/6NJ (B6NJ) genetic background generated using *Sox2-Cre* to ubiquitously delete exon 3. In agreement with other studies [14–16], *Sox2-Cre; Rasa3* null embryos displayed a phenotype indicative of severe hemorrhaging. Notably, significant pallor and decreased fetal liver size are also apparent in the mutant embryos compared to control (S2 Fig).

**Figure 1.**
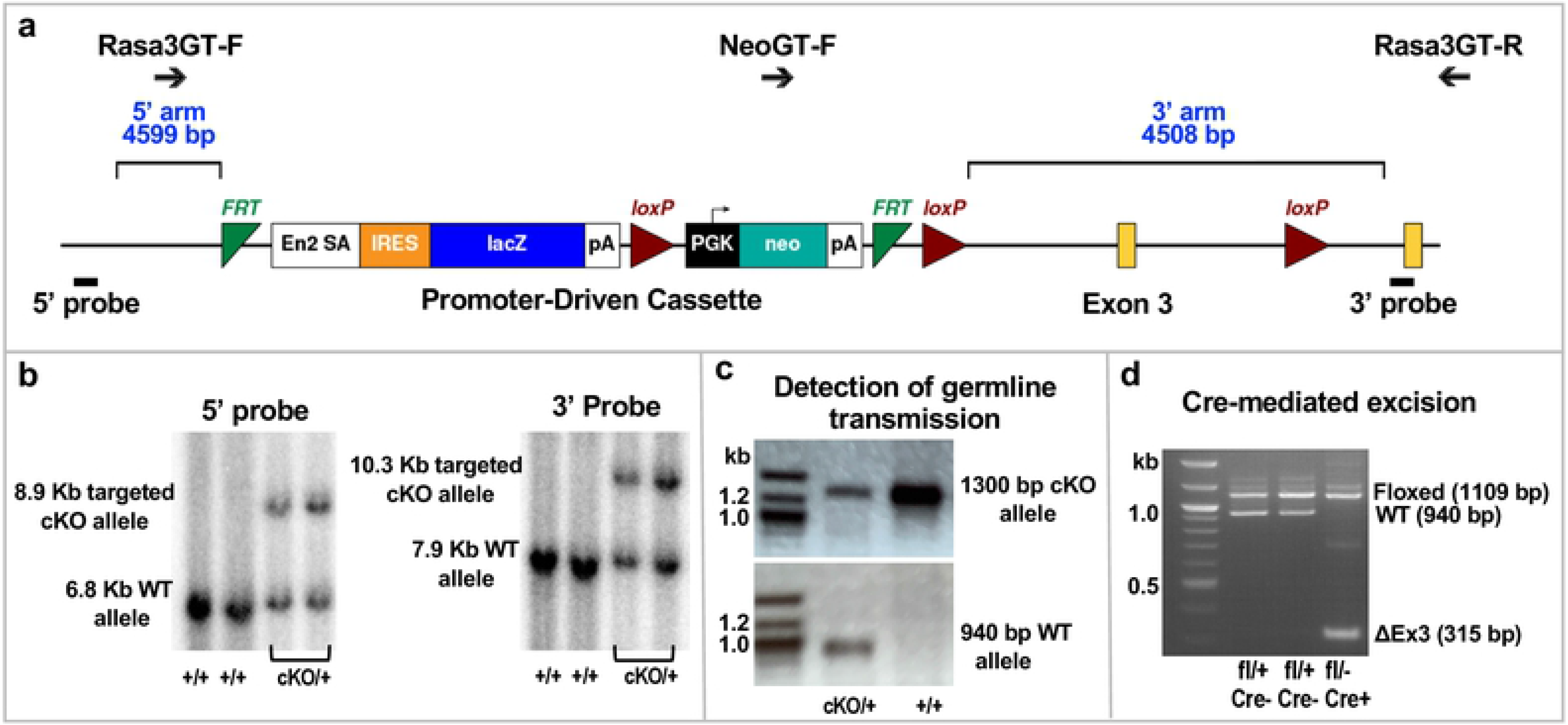
Generation of *Rasa3* conditional knockout (cKO) mice. **(A)** *Rasa3*^tm1a(KOMP)Wtsi^ knockout-first, promoter-driven conditional ready construct. Positions of genotyping (GT) primers, *FRT* (green) and *loxP* (red) sites, *Rasa3* exons (yellow), homology arms (blue) and flanking Southern blot probes (black rectangles) are shown. **(B)** Southern blots identifying correctly targeted cKO and wild type (WT) alleles in *BclI*- and *Bgl1*-digested ES cell genomic DNA using 5’ (left, *Bcl1* digest) and 3’ (right, *Bgl1* digest) flanking probes. **(C)** Detection of germline transmission using PCR to detect targeted cKO and WT alleles in genomic DNA. **(D)** PCR detection of *Rasa3* alleles following Flpe-mediated recombination to delete the promoter-driven cassette and Cre-mediated recombination to delete exon 3. Primer sequences for PCR reactions are provided in Supplementary Table 2.

### No abnormal phenotype is seen upon deletion of *Rasa3* in erythroid progenitors and precursors

We generated *Rasa3*^*fl*/−^ and *Rasa3*^*fl/fl*^ mice carrying *Epor-Cre* where *Cre* expression is driven by the erythropoietin receptor promoter beginning at the BFU-E stage and continuing through the proerythroblast stage of terminal erythroid differentiation [18]. Surprisingly, all *Epor-Cre; Rasa3* mutant offspring were normal in appearance at birth and at 6-8 weeks of age despite efficient deletion of exon 3 in red cell precursors (S3A Fig). Complete blood counts in adults did not differ from control in any parameters except for slightly decreased platelet counts and increased circulating reticulocytes (S3 Table), neither of which reached clinically significant levels nor approached the magnitude of changes seen in *scat* homozygotes (S4 Table). Spleen weight and peripheral blood morphology were normal as well in *Epor-Cre; Rasa3* mutant mice (S3B,C Fig). Western blotting of red cell membrane ghosts confirmed that abundant RASA3 protein was present in mutant red cell membranes (S3D Fig), suggesting that RASA3 produced prior to the BFU-E stage of erythroid development persists in terminally differentiated red cells.

### Deletion of *Rasa3* in hematopoietic stem and progenitor cells (HSPCs) and endothelial cells (ECs) recapitulates the null phenotype

To test the hypothesis that sufficient stable RASA3 protein is produced prior to the BFU-E stage of erythroid development to maintain terminally differentiating *Epor-Cre; Rasa3* mutant red cell precursors, we first examined *Rasa3* expression in normal B6J HSPCs. Indeed, expression of *Rasa3*, highest in early HSCs, declines by more than 80% in an enriched mixture [19] of BFU-E and CFU-E erythroid progenitors (Fig 2A). Examination of RNAseq data available online (nanexpression.mdibl.com [20]) reveals that *Rasa3* expression continues to decline as terminal differentiation proceeds (not shown). We next utilized *Vav-Cre* to delete *Rasa3* in HSPCs and ECs [21, 22]. No viable *Vav-Cre; Rasa3* mutant mice were recovered at birth; all died *in utero* at E12.5-13.5 with severe hemorrhage and significantly decreased fetal liver erythropoiesis (Fig 2B,C), precisely as observed in germline null *Sox2-Cre; Rasa3* mutants (S2 Fig). We obtained the same result when Cre expression was driven by Tie2 (*Tek-Cre,* Fig 2D,E), which is also expressed in both ECs and HSCs [22, 23].

**Figure 2.**
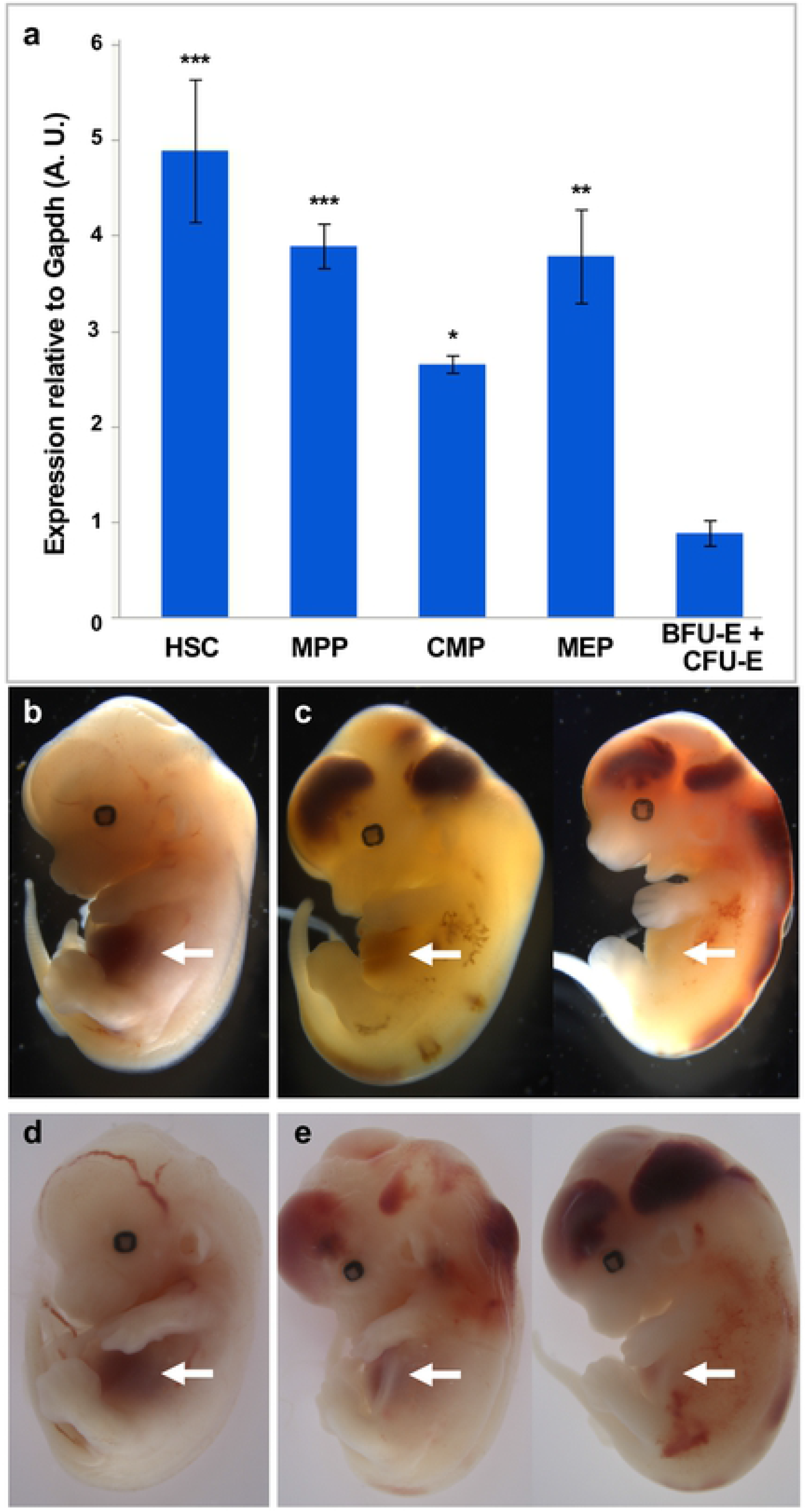
Deletion of *Rasa3* in hematopoietic and endothelial cells. **(A)** Expression of *Rasa3* in flow sorted adult B6J HSPCs (X ± SEM, n = 3). **(B)** E12.5-E13.5 *Vav-Cre*; *Rasa3* control and **(C)** mutant embryos. **(D)** E12.5-E13.5 *Tek-Cre; Rasa3* control and mutant **(E)** embryos. Mutant embryos recapitulate the germline null phenotype with severe hemorrhage and decreased fetal liver erythropoiesis (arrows). HSC, hematopoietic stem cells; MPP, multipotent progenitors; CMP, common myeloid progenitor; MEP, Myeloid-erythroid progenitor; BFU-E, burst forming unit-erythroid; CFU-E, colony forming unit-erythroid. *p < 0.05, **p < 0.01, ***p < 0.001 *vs*. BFU-E/CFU-E.

### Induced deletion of *Rasa3* in adults leads to severe anemia, leukopenia, and thrombocytopenia

We next utilized *Mx1-Cre* transgenic mice to delete RASA3 throughout the hematopoietic system in adult mice upon induction with polyinosinic:polycytidylic acid (Poly(I:C))(24, 25). Western blotting confirmed loss of RASA3 protein in *Mx1-Cre; Rasa3* mutant red cell membranes within 2 weeks of the final Poly(I:C) injection (Fig 3A) at which time the mice are profoundly anemic and thrombocytopenic with significant leukopenia (Fig 3B, Table 1).

**Fig. 3.**
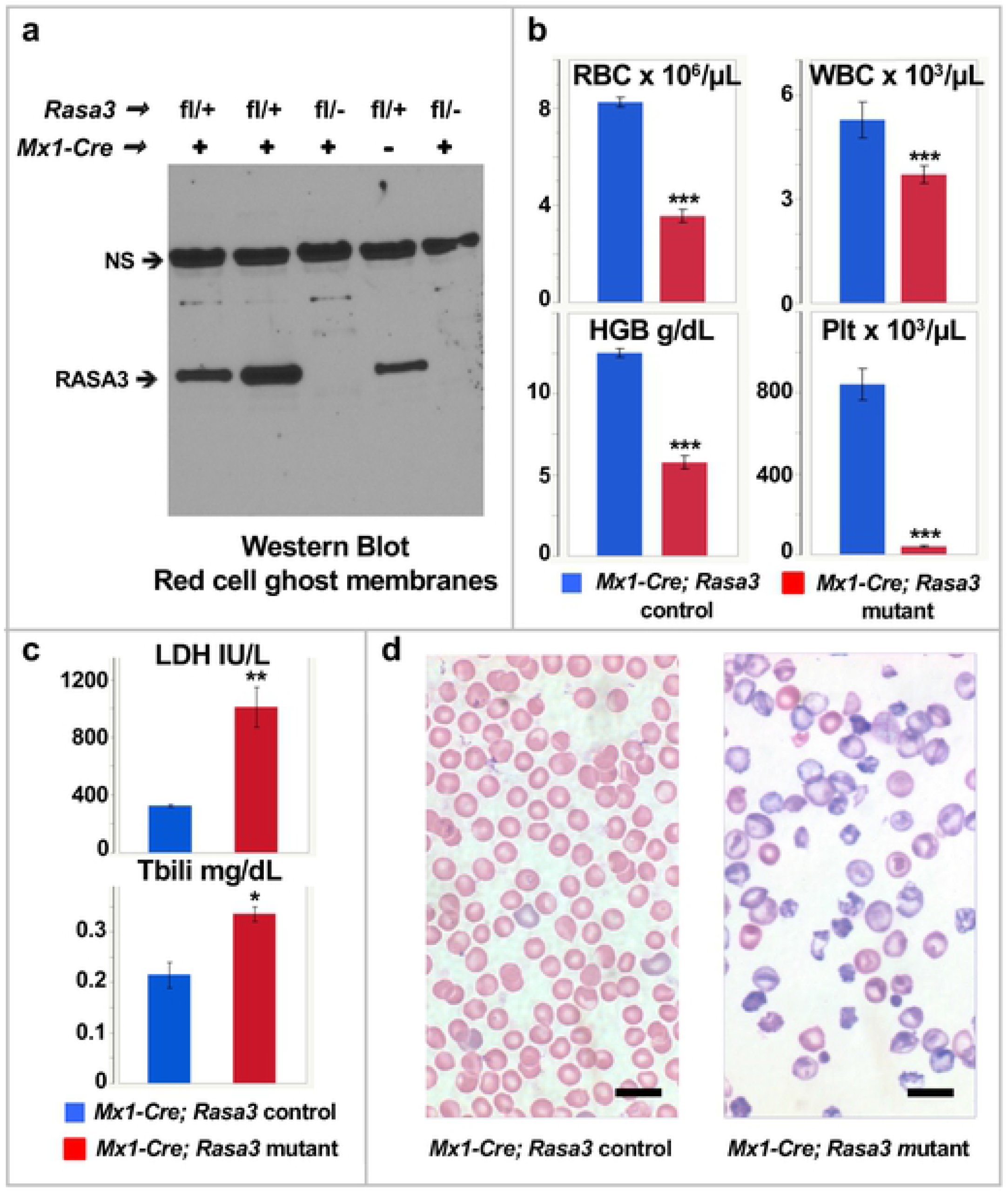
Deletion of *Rasa3* in adults leads to profound anemia. **(A)** Western blotting confirms loss of RASA3 protein in mutant red cell ghost membranes from pIpC treated *Mx1-Cre; Rasa3* adult mice. NS = non-specific band used as loading control. **(B)** Profound anemia and thrombocytopenia with significant leukopenia in pIpC treated *Mx1-Cre; Rasa3* mutant adults with **(C)** increased serum lactate dehydrogenase (LDH) and total bilirubin (Tbili) and **(D)** strikingly abnormal peripheral blood morphology. Values in b and c, X ± SEM for males only; complete CBC data is given for both sexes in Table 1. c, n = 4 (control) and 6 (mutant). Bar, 10 μM. *p < 0.05, **p < 0.01, ***p < 0.001.

**Table 1.**
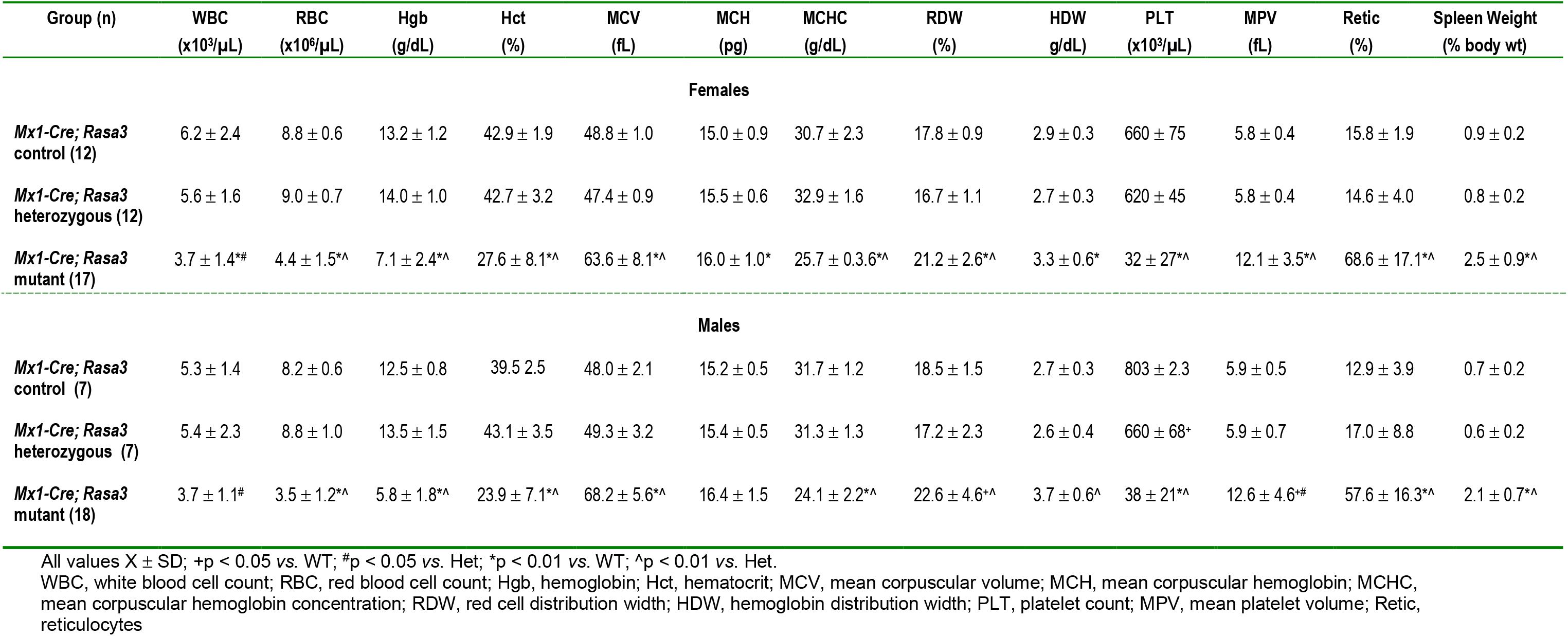
Complete blood counts in *Mx1-Cre; Rasa3* adult mice 6-8 weeks of age.

Spleen weight, percent circulating reticulocytes, mean corpuscular volume, and red cell and hemoglobin distribution widths are significantly increased in *Mx1-Cre; Rasa3* mutant mice compared to controls (Table 1). Serum LDH and total bilirubin, indicative of hemolysis, are increased (Fig 3C), and the peripheral blood morphology is markedly abnormal (Fig 3D). *Mx1-Cre; Rasa3* heterozygotes did not differ from control in any of these parameters (Table 1). Together, our studies using *Vav*-, *Tie2*- and *Mx1-Cre* confirm that production of RASA3 in HSPCs is required in order to maintain terminal erythroid differentiation and generate normal numbers of circulating mature erythroid cells as well as leukocytes and platelets.

### Bone marrow suppression in *Mx1-Cre; Rasa3* mutant mice

Basal erythropoiesis is suppressed in *Mx1-Cre; Rasa3* mutant mice; total, CD45^−^ erythroid, and CD45^+^ non-erythroid cell counts are all significantly depleted in mutant *vs.* control bone marrow (Fig 4A). Flow cytometric analyses using CD44, Ter119 and forward scatter (FSC) as markers of terminal erythroid differentiation [26] show that progression through terminal differentiation proceeds normally in *Mx1-Cre; Rasa3* bone marrow, keeping pace with both non-anemic controls and controls rendered anemic by phlebotomy (PHB), which do not differ from each other in any red cell parameters (Fig 4B,C; S5 Table). Thus, basal erythropoiesis is quantitatively suppressed but terminal differentiation *per se* is unaffected in the absence of RASA3.

**Figure 4.**
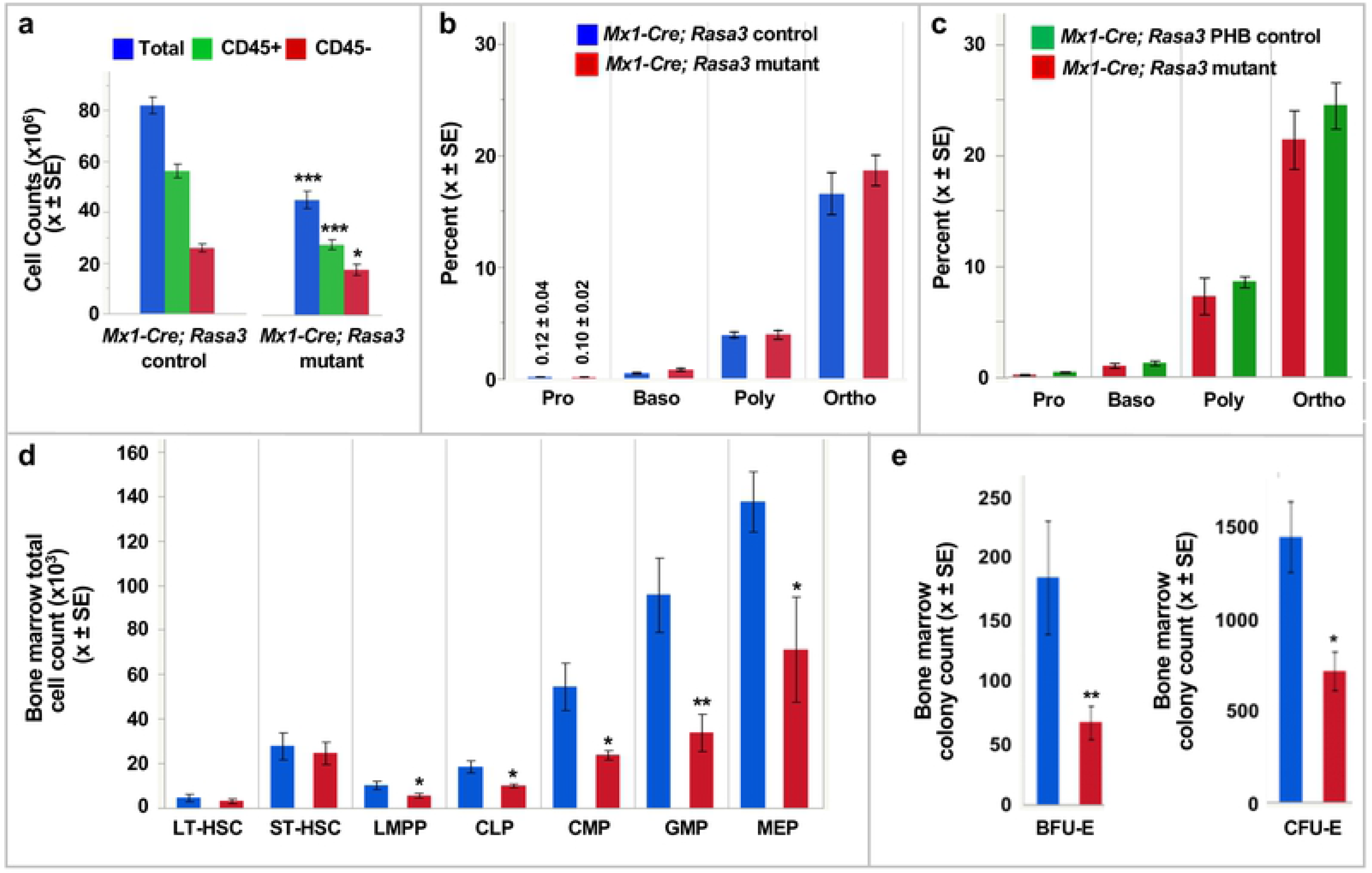
Suppression of bone marrow hematopoiesis in the absence of RASA3. **(A)** Total, CD45− erythroid, and CD45+ non-erythroid cell counts are significantly depleted in *Mx1-Cre; Rasa3* mutant bone marrow. n = 8 **(B,C)** Terminal erythroid differentiation progresses normally in mutant bone marrow compared to both non-anemic (n = 8) and phlebotomized anemic (n = 6) controls. **(D)** Total counts of all hematopoietic progenitor cell types are significantly decreased in *Mx1-Cre; Rasa3* mutant bone marrow *vs*. control (n= 8). LT- and ST-HSCs do not significantly differ. **(E)** Decreased BFU-E and CFU-E colony formation capacity in mutant bone marrow. n = 3. Equal numbers of mutant and control were plated in colony forming assays. LT-HSC, long term hematopoietic stem cell; ST-HSC, short term hematopoietic stem cell; LMPP, lymphoid-primed multipotent progenitor; CLP, common lymphoid progenitor; CMP, common myeloid progenitor; GMP, granulocyte monocyte progenitor; MEP, Myeloid-erythroid progenitor. All values X ± SEM. *p < 0.05, **p < 0.01, ***p < 0.001.

We next determined the status of hematopoiesis at the level of HSPCs using flow cytometric approaches. The frequencies of all HSPC populations in *Mx1-Cre; Rasa3* mutant and control bone marrow are the same (S4A Fig) as is the absolute number of LT- and ST-HSCs (Fig 4D). However, the absolute number of all progenitors is significantly decreased in mutant bone marrow compared to control (Fig 4D). Moreover, in keeping with reduced progenitors including megakaryocyte-erythroid progenitors (MEPs), the functional capacity of mutant bone marrow to produce BFU-E and CFU-E colonies *in vitro* is strikingly reduced (Fig 4E), all of which would be predicted to contribute to pancytopenia in the absence of RASA3.

### Stress erythropoiesis in RASA3 mutant spleen

Spleen erythropoiesis increases significantly under anemic stress [27, 28]. In *Mx1-Cre; Rasa3* mutant mice, stress erythropoiesis is established within two weeks of induction of anemia; the spleen is grossly enlarged (Table 1) and its normal nodular architecture is effaced by expansion of the red pulp (S5 Fig). Flow cytometric analysis reveals that the total cell number is increased in mutant spleen compared to control, although it fell just short of statistical significance (p = 0.0531). The increase in total cell number reflects expansion of the erythroid compartment, as CD45^−^ erythroid but not CD45^+^ non-erythroid cells are significantly increased compared with controls (Fig 5A).

**Figure 5.**
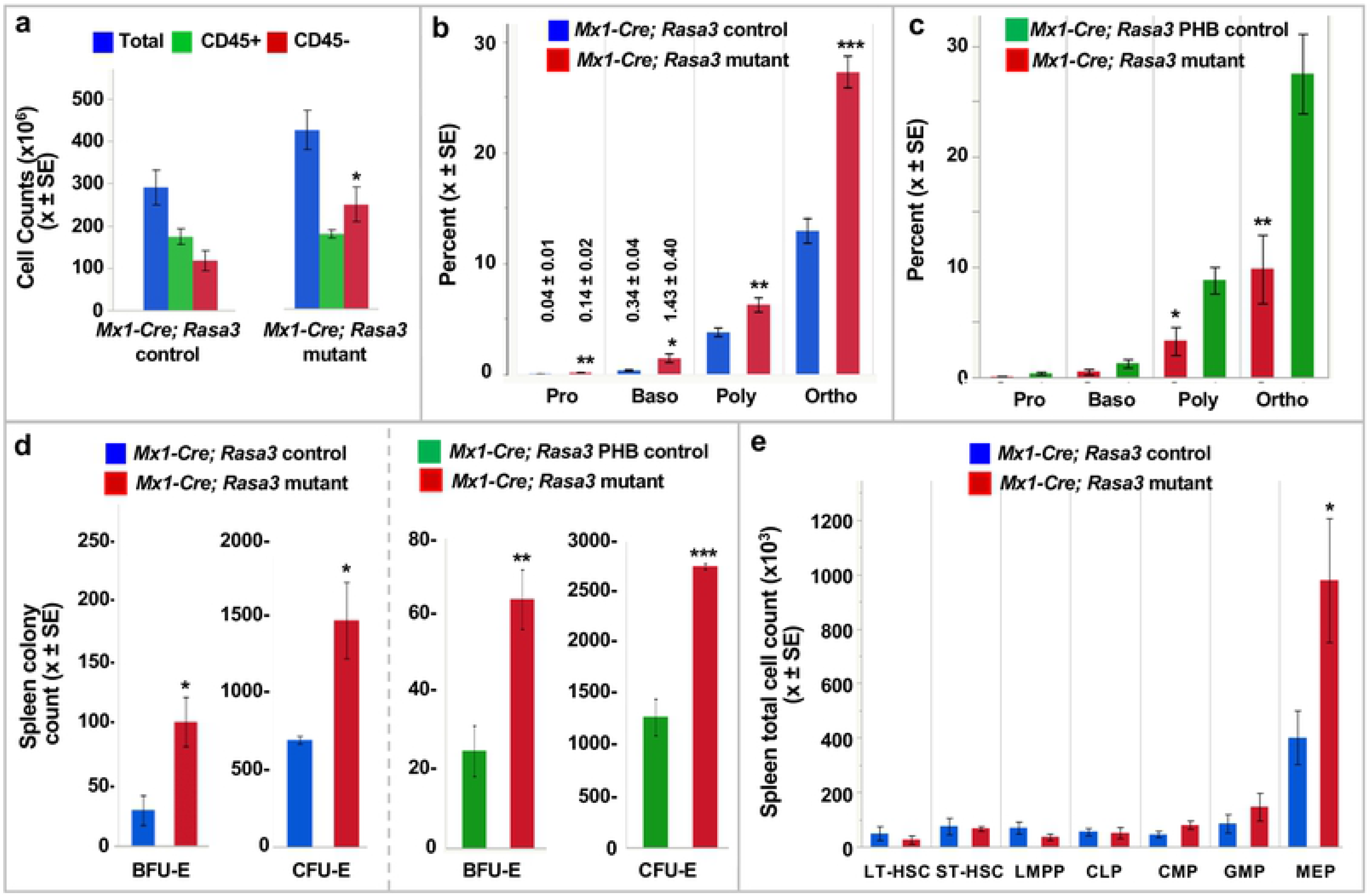
Blunted spleen stress erythropoiesis in the absence of RASA3. **(A)** Total CD45− erythroid cells are significantly increased in *Mx1-Cre; Rasa3* mutant adult spleen. n = 8. **(B)** Terminal differentiation in the mutant spleen is accelerated compared to *Mx1-Cre* negative control mice, showing increased precursors at all stages. n = 8. **(C)** Stress erythropoiesis in *Mx1-Cre; Rasa3* mutant spleen is dramatically blunted compared to *Mx1-Cre* control mice rendered anemic by phlebotomy (PHB). n = 6. **(D)** Increased BFU-E and CFU-E colony formation in mutant spleen. n = 3. Equal numbers of mutant and control were plated in colony forming assays. **(E)** Megakaryocyte-erythroid progenitors are increased in number in *Mx1-Cre; Rasa3* mutant spleen; no other significant differences are seen. n = 8. LT-HSC, long term hematopoietic stem cell; ST-HSC, short term hematopoietic stem cell; LMPP, lymphoid-primed multipotent progenitor; CLP, common lymphoid progenitor; CMP, common myeloid progenitor; GMP, granulocyte monocyte progenitor; MEP, Myeloid-erythroid progenitor. All values X ± SEM. *p < 0.05, **p < 0.01, ***p < 0.001.

Flow cytometric analyses of terminal erythroid differentiation reveal an increase in all precursor stages from pro- to orthochromatic erythroblasts in *Mx1-Cre; Rasa3* mutant compared to control spleen (Fig 5B). The spleen erythropoietic response in *Mx1-Cre; Rasa3* PHB anemic controls, however, was substantially higher than in *Mx1-Cre; Rasa3* mutants (Fig 5C). Thus, while comparison to control mice suggests that a stress response is initiated in *Mx1-Cre; Rasa3* mutant spleen, comparison to PHB control mice reveals that the response is strikingly less robust than that seen in anemic mice in which the *Rasa3* locus is intact. Moreover, since the reticulocyte response from phlebotomy originates from the precursor pool [29–31], these data suggest that the stress erythropoietic response in *Mx1-Cre; Rasa3* mutants is initiated at the progenitor level. Colony-forming assays on *Rasa3* mutant, anemic PHB controls, and non-anemic controls support this hypothesis. BFU-E and CFU-E colony numbers are significantly increased in *Rasa3* mutant spleen compared to both non-anemic and anemic (PHB) controls, which are roughly comparable to each other (Fig 5D). Moreover, MEPs are increased both in number and in frequency in *Mx1-Cre; Rasa3* mutant spleen compared to control, although no other significant differences are seen in HSPC populations (Fig 5E, S4B Fig). Together, our results support the hypothesis that although deletion of *Rasa3* leads to an increase of early erythroid progenitors in the spleen, both in number and function, enhanced erythropoiesis is not propagated effectively through terminal differentiation.

### Genetic background profoundly influences the *Rasa3* mutant phenotype

Dramatic phenotypic differences are observed in engineered germline *Rasa3* null mice on the B6J genetic background compared to *scat* mice carrying a missense mutation on BALB/cBy (cBy). While a subset of cBy-*scat* mice dies *in utero*, 10-15% are born and show a characteristic pale, bruised phenotype as neonates (Fig 6A). *Rasa3* germline null mice on either a mixed B6,129 (Fig 2) or a pure inbred B6NJ (S2 Fig) genetic background are 100% embryonic lethal. To examine the effects of genetic background more closely, we first examined complete blood counts in B6NJ adults; B6NJ *Mx1-Cre; Rasa3* null mice display severe anemia, thrombocytopenia and leukopenia, closely mirroring the B6;129 *Mx1-Cre; Rasa3* null phenotype (S6 Table). We next transferred the *scat* missense allele to the B6J background to create a fully congenic line, B6J.cBy-*scat*. In this case, a dramatically different phenotype emerged in B6J congenic *vs*. cBy *scat* m*ice* despite both carrying the same missense allele. No affected offspring showing the easily recognizable *scat* phenotype at birth were identified in B6J.cBy-*scat*/+ intercrosses at any outcross generation. Genotyping of 154 neonates at the most recent outcross generation (N = 21) revealed 106 (68.8%) *scat*/+ and 48 (31.2%) wildtype mice; no *scat/scat* mice were detected, confirming 100% *in utero* lethality. Examination and genotyping of 64 congenic fetuses at E12.5-E14.5 revealed the expected 25% *scat* homozygotes; all exhibited evidence of severe bleeding with strikingly pale fetal livers (Fig 6B). Thus, congenic B6J.cBy-*scat* homozygotes differ markedly from cBy-*scat/scat* and mimic germline B6J/B6NJ null mutations, confirming a striking effect of genetic background.

**Figure 6.**
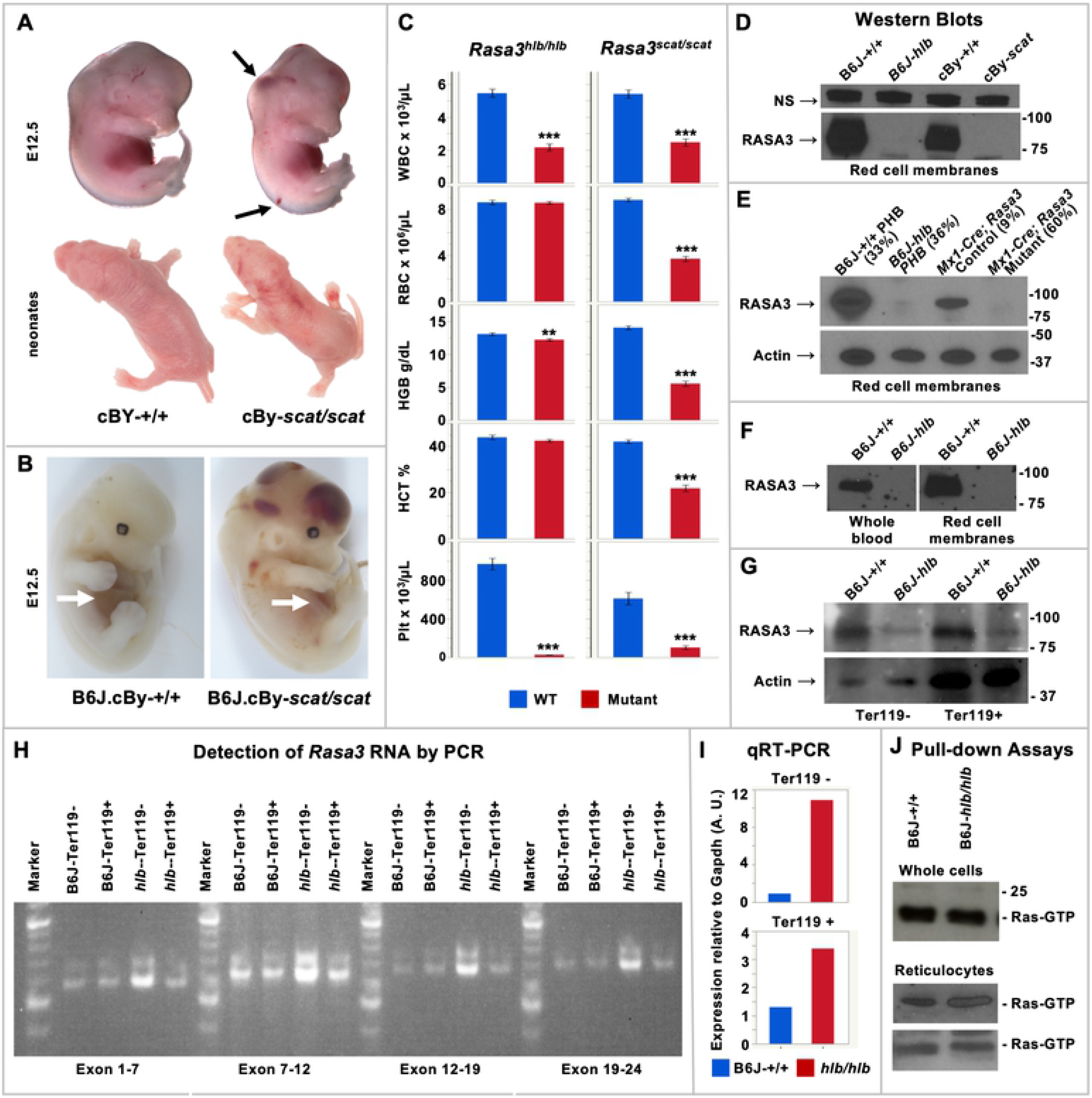
Effects of genetic background and molecular variant on RASA3 mutant phenotype. **(A)** E12.5 (top) and neonatal (bottom) BALB/cBy (cBy) control and *scat/scat* mice. Pallor and extensive bruising are consistently observed in newborn *scat* homozygotes. **(B)** Congenic B6J.cBy *scat* homozygote displaying extensive hemorrhaging and a strikingly pale fetal liver *vs*. control (arrows). Examination of multiple litters from E12.5 to E14.5 showed no live B6J-*scat/scat* mice. **(C)** *Hlb381* homozygotes (*hlb/hlb*) carrying missense mutation H794L are leukopenic and thrombocytopenic but differ markedly from *scat/scat* mice (G125V) in that severe anemia is absent. Supplemental Tables S4 and S7 provide complete blood count data for *scat* and *hlb381*, respectively, at 3-4 weeks of age. All values X ± SEM. **p <0.01, ***p < 0.001 *vs*. strain specific WT control. **(D-G)** RASA3 Western blots: **(D)** control and homozygous mutant *hlb* and *scat* red cell membranes (NS = non specific band serving as loading control from same blot), **(E)** phlebotomized (PHB) control and mutant *hlb* and *Mx1-Cre* red cell membranes (number in parentheses = circulating reticulocyte percentage), **(F)** control and *hlb* whole blood and red cell membranes, and **(G)** purified control and *hlb* Ter119− and Ter119+ spleen cells. **(H)** PCR detecting cDNA across all exons in control and *hlb* Ter119− and Ter119+ spleen cell mRNA. **(I)** qRT-PCR reveals increased *Rasa3* expression in *hlb* Ter119− and Ter119+ spleen cells compared to control. **(J)**. Pull down assays reveal normal levels of active Ras-GTP in *hlb* whole blood and reticulocytes.

### Dramatic effects of molecular variant on disease phenotype

Previously, we identified the recessive N-ethyl-N-nitrosourea (ENU) induced missense mutation in *Rasa3* (H794L) in a forward genetic mutagenesis screen in B6J mice [14, 32]. The mutations lie in the C-terminal 80 residue region of RASA3 that does not show homology to any known protein domains. Phenotypically the mutant model, designated *hlb381*, differs dramatically from *Rasa3* germline null, cBy-*scat* and congenic B6J.cBy-*scat* homozygotes. *Hlb381* homozygotes are born at the expected Mendelian ratio and survive normally. No pallor or evidence of bruising has ever been noted at birth or postnatally. Complete blood counts reveal severe thrombocytopenia and leukopenia at birth and throughout life. Unlike cBy-*scat* (G125V), however, severe anemia is not present, although some evidence of a mild compensated anemia (increased spleen weight and circulating reticulocyte percentage) is seen at 3-4 weeks of age (Fig 6C, S7 Table), most evident in males, that is largely resolved by 6 weeks of age (S8 Table). The spleen architecture is normal, with clearly delineated white and red pulp areas (S6 Fig). Thus, different *Rasa3* missense alleles, G125V (B6J.cBy-*scat*) and H794L (B6J-*hlb381*), confer dramatically different phenotypes on the same (B6J) genetic background, adding further data emphasizing the significant structure-function differences between RASA3 amino acid variants.

Membrane localization of RASA3 is required for GAP activity to downregulate RAS and RAP activity [1, 4]. RAS, but not RAP, is present in mouse erythrocytes; RAP is absent in mouse red cells but abundant in platelets[(12, 14, 33]. Loss of RASA3 activity would be predicted to increase active GTP-bound RAS and RAP. Indeed, our previous studies in *hlb381* revealed increased RAP-GTP in platelets, leading to spontaneous platelet activation and markedly increased clearance of circulating platelets [14]. We also showed previously that active RAS-GTP is increased in RASA3 mutant *scat* reticulocytes as a result of mislocalization of RASA3 to the cytosol leading to abnormal erythropoiesis [12]. The lack of severe anemia in *hlb381* leads to the prediction that, unlike G125V in *scat*, H794L does not affect membrane localization or RAS activation. To test this hypothesis, we performed western blots and pull-down assays. Surprisingly, although easily detectable in B6J controls, RASA3 is undetectable in *hlb381* red cell membranes even upon over-exposure of ECL western films (Fig 6D). RASA3 is most abundant in reticulocytes, but it declines rapidly via the exosome pathway during reticulocyte maturation [12]. Despite this, even when significant reticulocytosis is induced via phlebotomy, RASA3 remains undetectable in *hlb381* red cell membranes (Fig 6E) and whole cells (Fig 6F). Only when we examined purified early erythroid cells from spleen, were we able to detect RASA3 protein in *hlb381* albeit at much lower levels than controls (Fig 6G). Notably, prior studies in platelets showed that RASA3 protein is also reduced in homozygous *hlb381* platelet lysates to levels comparable to those in platelets from heterozygous mice carrying a RASA3 null allele, suggesting that the H794L allele markedly impairs RNA expression or RASA3 protein stability [14]. Our findings here confirm instability of the H794L RASA3 allele, as conventional PCR analysis of spleen erythroid cells detects cDNA for all *Rasa3* exons (Fig 6H). Moreover, significantly more PCR product is seen for *hlb381* phlebotomized spleen erythroid cells, particularly Ter119− progenitors, compared to WT; qRT-PCR confirms increased *Rasa3* expression in *hlb381* homozygotes (Fig 6I).

Finally, in contrast to previous studies showing increased RAP-GTP in *hlb381* platelets [14], pull-down assays failed to show increased active RAS-GTP in *hlb381* erythroid cells consistently, notably in purified reticulocyte fraction (Fig 6J). Together, these data suggest that mislocalization of RASA3 from the membrane to the cytosol with resulting increased active RAS-GTP leading to abnormal erythroid differentiation, as occurs in *scat*, does not occur in *hlb381* and could explain why significant anemia is absent in *hlb381*.

### Analyses of global transcriptomes in *scat* suggest a primary role of spleen in disease genesis and differentiates potential mechanisms underlying crisis *vs.* partial remission

To determine global transcriptome changes in RASA3 defective hematopoietic tissues and cells, we performed RNAseq using 3-5 biological replicates of whole spleen and bone marrow and flow-sorted HSPC populations from *scat* homozygotes and their WT littermates (S9 Table). Furthermore, the cyclic nature of *scat* disease progression, in which rare full remissions and, more commonly, partial remissions occur (Fig 7A) allowed comparison of the same tissue/cell populations from *scat* crisis (cr) and *scat* partial remission (pr) mice. Mice were classified as pr based on erythroid parameters intermediate to WT and cr with the exception of the reticulocyte percentage and spleen weight, which are expected to remain high during recovery from anemia. Complete blood counts for mice in each group used for RNAseq are given in S10 Table. Flow-sorted populations obtained were megakaryocyte-erythroid progenitors (MEP) and so-called “stem and myeloid progenitors” (SMP). Because Sca1 (*Ly6a*) is expressed at very low levels in BALB mice(34), HSCs cannot be separated reliably from common myeloid progenitors. Therefore, the Lin^−^Sca^−^Kit^+^CD34^+^ CD16/32^lo^ SMP population was used for these studies [35].

**Figure 7.**
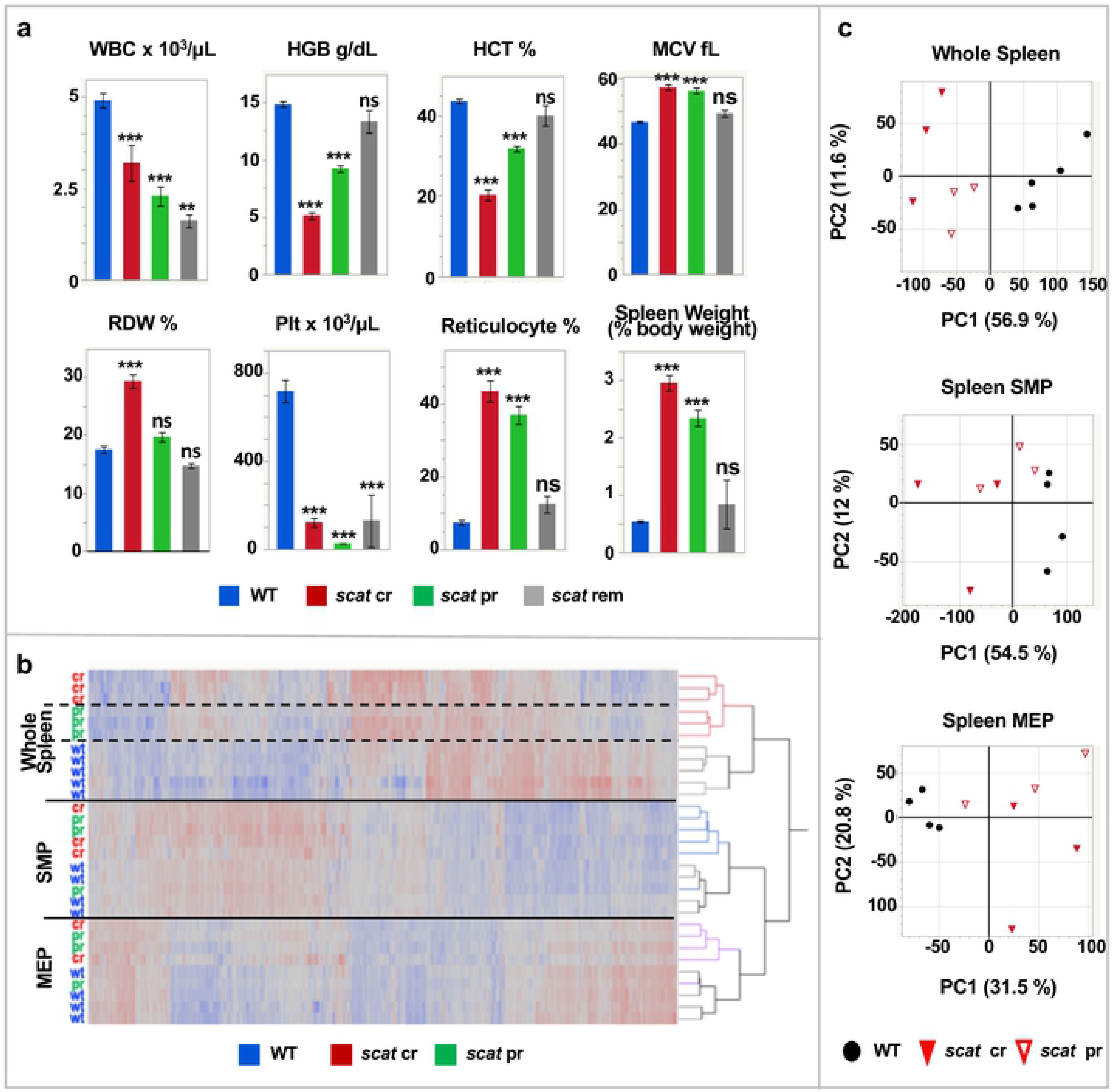
Transcriptome analyses in *scat* mice. **(A)** Blood counts in control (WT, n = 38), *scat* crisis (cr, n = 25), *scat* partial remission (pr, n = 22) and *scat* full remission (rem, n = 4) and spleen weights (n = 24, 19, 12, and 2, respectively). Data are male and female combined. All values X ± SEM. **p < 0.01, ***p < 0.001, ns not significant. **(B)** Hierarchal clustering and **(C)** principle component analysis of expression differences clearly separates whole spleen, SMP, and MEP samples (solid lines) and WT, cr and pr within whole spleen (dashed lines).

Differentially expressed gene (DEG) lists are given in S12-S24 Tables. Hierarchal clustering of DEGs in spleen and spleen-derived erythroid HSPCs clearly separates whole organ, SMP, and MEP populations (Fig 7B). Moreover, in whole spleen there is a clear clustering of *scat* phenotypes (cr, pr, wt) that is not maintained in SMP and MEP cells (Fig 7B). Principle component analysis (PCA) reveals that the first principle component (PC1) dominates and segregates by genotype (*scat* vs. WT) in all spleen-derived samples. In whole spleen PC1 additionally segregates cr from pr suggesting that functionally distinct mechanisms are at play among the three phenotypes in the spleen. (Fig 7C). In bone marrow, PC1 separates *scat* from WT in whole bone marrow only and does not distinguish cr and pr (S7 Fig).

To explore functions associated with altered expression patterns we performed gene ontology (GO) analysis using DAVID [36]. Clearly, many comparisons of expression differences are possible between the many datasets generated; here, we present those we found to be most informative. Additional comparisons can made using DEG lists provided (S12- S24 Tables). In whole spleen, when DEGs for all mutants (cr and pr combined) are compared to WT controls using GO terms for Biological Process, terms associated with cell cycle and immune function predominate; cell cycle terms dominate genes upregulated in mutants, and immune function down-regulated (S8 Fig). Notably, cell cycle [37] and immune cell changes [33] have been documented in *scat*. Similar comparisons of bone marrow DEGs are dominated by immune and chromatin-associated terms (S8 Fig). Analysis of KEGG pathways in DAVID were informative in spleen SMP and MEP (Fig 8A). In SMP DEGs, the arrhythmogenic right ventricular cardiomyopathy (ARVC) pathway is significant. A closer look at the DEGs associated with this pathway reveals genes encoding components of adherens junctions (AJ) and desmosomes.

**Figure 8.**
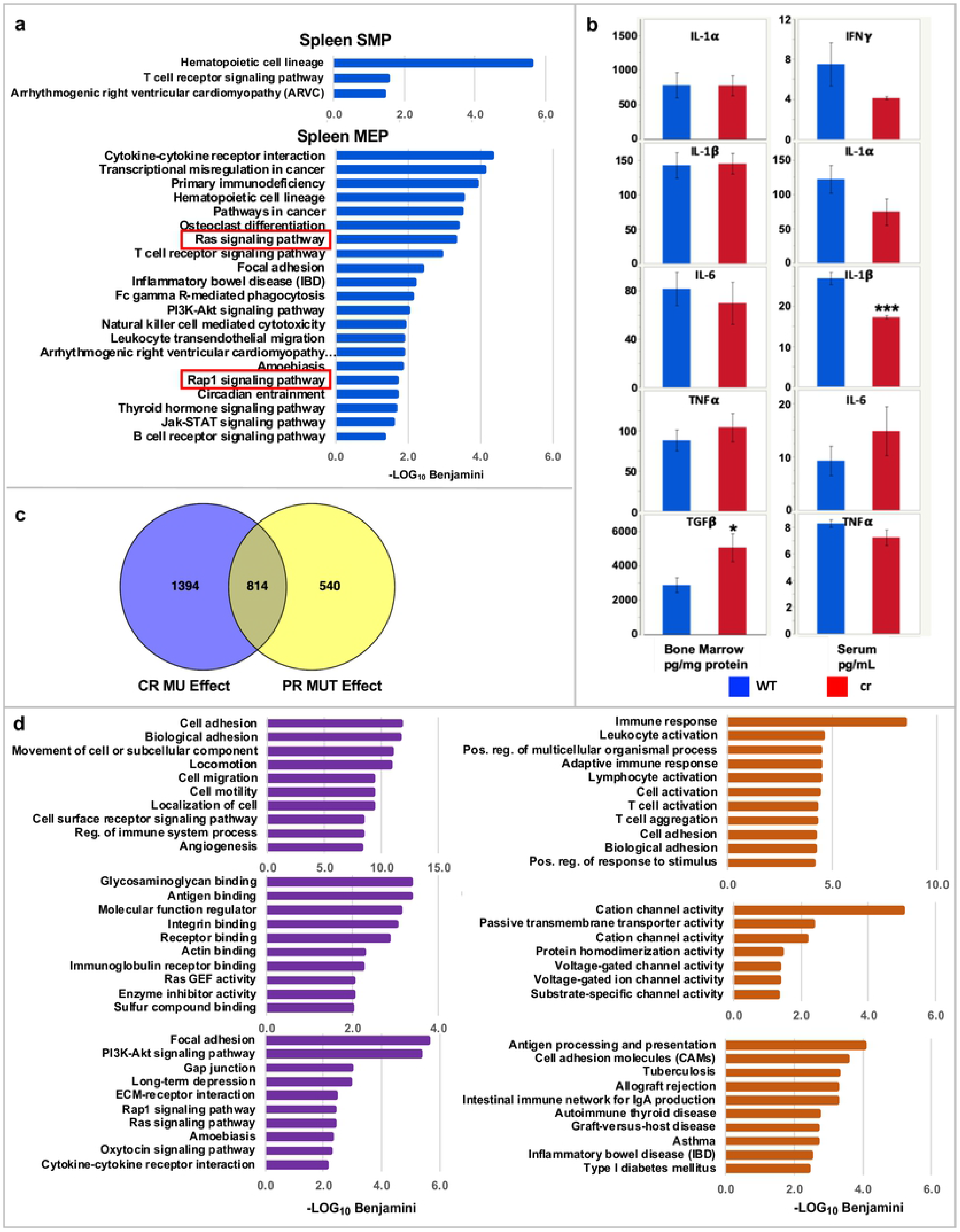
Gene ontology analyses of differentially expressed genes. **(A)** Significant pathways (KEGG) associated with spleen SMP and MEP differential expression. **(B)** Cytokine levels in bone marrow fluid and serum. **(C)** Venn diagram of expression differences in cr *vs*. pr. **(D)** Significant biological process (top), molecular function (middle) and KEGG pathways (bottom) exclusive to cr (left, purple) and pr (right, orange) DEGs.

Notably, the original description of RASA3 knockout noted underdeveloped AJs between capillary endothelial cells [15]. In similar analyses of spleen MEP DEGs, RAS and RAP signaling were among significant pathways (Fig 8A). Cytokine interactions also emerged. A preliminary survey of a small subset of cytokines in bone marrow fluid showed increased TGFβ and, in serum, decreased IL-1β (Fig 8B).

Given that mechanisms known to be associated with the *scat* disease phenotype emerged in the above analyses, we were encouraged that comparison of cr vs pr could identify processes associated with recovery of anemia. We analyzed differential expression in cr vs. pr across all samples combined to determine an overall mutation effect; 540 DEGs were unique to pr, and 1384 to cr (Fig 8C) [38]. Significant GO terms for Biological Process, Molecular Function, and KEGG pathways unique to cr and pr are shown in Fig 8D. Importantly, examination of these terms reveal that those exclusive to cr “fit” our expectations to a large degree given what is known about the *scat* crisis phenotype from this and prior studies[(12, 14]. Terms and processes exclusive to pr provide some surprising, hypothesis-driving new directions for future studies. Notably, the most significant Molecular Function GO terms that emerged are all related to channel activity, suggesting that changes in red cell transporter activity are central to partial recovery of the *scat* anemia.

## Discussion

Studies in the *scat* mutant mouse model, characterized by severe anemia, thrombocytopenia and leukopenia, and confirmed in zebrafish by morpholino knockdown firmly established a critical non-redundant role of RASA3 in vertebrate blood formation [12]). The original gene knockout study in which a catalytically inactive form of RASA3 was produced [15] showed that germline loss of RASA3 activity results in extensive hemorrhage, a phenotype that was attributed to quantitatively decreased adherens junctions between endothelial cells. Subsequent studies confirmed that the RASA3-RAP axis is critical in endothelial cell adhesion and vessel development [16]. *Scat* mutants carrying the G125V missense mutation also show evidence of hemorrhage, although less severe, *in utero* (Fig 6A), which may account for the fact that *scat*/+ intercrosses produce 10-15% homozygous *scat* newborns *vs.* the expected 25%. Postnatally, cranial hemorrhage frequently develops during crisis episodes in *scat* [12, 13]. Interestingly, vascular malformations in the brain accompanied by hemorrhagic stroke were described recently in mice expressing activated HRAS alleles [39]. Whether such a mechanism underlies cranial bleeding in *scat* awaits further study.

Here we show that *Rasa3* null embryos not only show massive bleeding but evidence of insufficient fetal liver erythropoiesis as well. Significantly, this phenotype is also observed in conditional *Vav-Cre*; *Rasa3* mutant embryos, indicating a requirement for RASA3 in the embryo at the hemangioblast stage of vascular/hematopoietic development. To investigate hematopoiesis further, we first focused on terminal erythroid differentiation, as a pronounced block at the poly- and orthochromatophilic stages was previously documented in *scat* [12]. Surprisingly, no phenotype emerged in *Epor-Cre; Rasa3* mutants, and RASA3 protein persisted in erythroid cells suggesting that RASA3 produced in progenitors maintained erythroid terminal differentiation. Indeed, when *Rasa3* is deleted in adults using *Mx1-Cre*, the severe, pancytopenic *scat* phenotype emerges. Bone marrow hematopoiesis is clearly suppressed; total cellularity is decreased, all progenitors are significantly decreased in number, and the functional capacity to produce erythroid progenitors (BFU-E, CFU-E) is significantly decreased (Fig 4).

In addition to suppressed bone marrow function in *Mx1-Cre*; *Rasa3* mutants, stress erythropoiesis in the spleen is ineffective, as has been described previously in *scat* mice [12]. Although increased numbers of all red cell precursors are produced in the *Mx1-Cre*; *Rasa3* mutant spleen compared to non-anemic controls, their numbers fall far short of those produced in PHB anemic control mice in which RASA3 is intact. Notably, unlike the *scat* model, terminal erythropoiesis is not delayed at any stage in *Mx1-Cre*; *Rasa3* mutants. However, the models differ in many respects including their ages and the nature of the RASA3 defect, as well as the disease itself, i.e., chronic vs. acute. Although terminal erythroid differentiation in *Mx1-Cre*; *Rasa3* mutant spleen falls short of PHB anemic controls, BFU-E and CFU-E colony forming ability is significantly greater (Fig 5) than both anemic and non-anemic controls. Thus, the *Mx1-Cre*; *Rasa3* mutant spleen initiates a stress response at the progenitor level that fails to be propagated through the precursor stages where rapid amplification of red cell production would occur.

The phenotypic differences previously described amongst various RASA3 mouse models are striking [12, 14, 16, 17, 40]. To address this aspect of RASA3 function, we repeatedly backcrossed the *scat* mutation onto the B6J genetic background, producing a fully congenic strain. Remarkably, 100% of B6J-*scat/scat* congenic mice die *in utero* (E12.5-13.5) recapitulating the germline null phenotype. Thus, genetic modifiers differing between BALB and B6J dramatically alter the effects of the RASA3 G125V mutation. The molecular variant also has a profound impact, as exemplified by the chemically induced *hlb381* model carrying a missense mutation, H794L, near the C-terminus of RASA3 in a region that is not structurally related to any known protein domains. *Hlb381* is on the B6J genetic background but survives normally, unlike B6J congenic *scat* mice. Previously, we showed that RASA3 protein levels were decreased in *hlb381* platelets, suggesting that the mutant protein was unstable. We also showed that as a consequence of decreased RASA3, active RAP-GTP levels increased in *hlb381* platelets [14]. Here, despite significantly increased mRNA expression, RASA3 protein is not detected in unfractionated circulating *hlb381* whole cells, red cell membrane ghosts or purified reticulocytes, but is detected at much reduced levels in purified spleen Ter119^+^ and Ter119^−^ cells, confirming instability of RASA3 H794L (Fig 6).

Decreased stability of RASA3 H794L and failure to bind to the membrane in reticulocytes and mature red cells would suggest increased active RAS-GTP with its consequent aberrant influence on erythropoiesis would arise in *hlb381* mice, as has been described in mutations in which RAS is constitutively activated, including *scat*. However, we failed to show consistently increased active RAS-GTP in *hlb381* erythroid cells, indicating that H794L does not significantly alter RAS activation and perhaps accounting, at least in part, for the absence of significant anemia in *hlb381*. The data in total suggest that mechanisms in addition to failure of mutant RASA3 to bind the membrane in *scat* cells and/or currently unknown compensatory mechanisms and genetic modifiers operating in *hlb381* also contribute to influence hematopoiesis in mouse models of *Rasa3* defects. The data further suggest that G125V alters RASA3’s GAP activity for both RAS and RAP, while H794L primarily affects RAP activation.

Of the RASA3 deficient models described to date, only *scat* shows a variable phenotypic progression of crisis (cr) and partial remission (pr). We exploited this aspect of the *scat* disease progression to examine global transcriptome differences between WT, cr, and pr HSPCs (SMP, MEP) and whole tissues (spleen and bone marrow). Remarkably, analyses of differential expression suggest a primary influence of the spleen on disease progression. Hierarchal clustering separated expression differences by cell/tissue type derived from both spleen and bone marrow, but only in whole spleen did the three phenotypes (WT, cr, pr) partition into distinct clusters (Fig 7). Principle component analysis confirmed a striking phenotype effect on differential expression in the spleen (Fig 7). Analysis of functional annotation GO terms unique to pr reveals unexpected differences compared to cr in that the most significant terms all related to channel activity (Fig 8), suggesting a novel direction for further study.

## Methods

### Mice

All mice were maintained in climate- and light cycle-controlled animal facilities at The Jackson Laboratory. Acidified water and 5K52 chow (PMI LabDiet) were provided *ad libitum*. The spontaneous co-isogenic *scat* mutation arose originally on the inbred BALB/cBy substrain. Subsequently, *scat* was outbred with the BALB/cByJ substrain for three generations and then propagated by brother-sister mating in the research colonies of Dr. Luanne Peters at The Jackson Laboratory. Hence, the official designation of the *scat* mutant strain is CByJ;CByRasa3^*scat*^/LLP. All mouse strains used in this study are given in Table S1.

All experiments were performed in accordance with National Institutes of Health Laboratory Animal Care Guidelines and were approved by the Animal Care and Use Committee (ACUC) of The Jackson Laboratory.

### Conditional knockout (cKO) strains

We generated a conditional targeted allele for *Rasa3* using the *Rasa3*^tm1a(KOMP)Wtsi^ Knockout-first, promoter-driven conditional ready construct (Fig 1) generated by the trans-NIH Knock-Out Mouse Project (KOMP) and obtained from the KOMP Repository (www.komp.org). The construct was electroporated into 129S1/SvImJ (JAX Stock# 002448)-derived ES cells and cultured with G418 selection using standard protocols. Genomic DNA from ES cell clones was digested with *Bcl1* and *Bgl1* to assess integrity of the 5- and 3-prime homologous recombination sites, respectively, by Southern blotting using 738 bp 5-prime and 649 bp 3-prime PCR-generated flanking probes. Primer sequences are provided in S2 Table. Blastocyst (C57BL/6J) injection and embryo transfer were performed using standard techniques. Male chimeras were mated to C57BL/6J mice to generate heterozygous *Rasa3*^*cKO*/+^ mice on the hybrid B6;129S1 genetic background. *Rasa3* and *Cre* genotypes in newborn progeny were determined initially using PCR (S2 Table) and subsequently by real time PCR with specific probes designed by Transnetyx.

Targeted mice using the same construct were subsequently generated by the Jackson Laboratory KOMP Production Center on the inbred C57BL/6NJ (B6NJ) genetic background [41].

### Breeding

The general breeding strategy is shown diagrammatically in Fig S1. Mice carrying the targeted, cKO allele were crossed to *ACT-Flpe* mice to excise the IRES:*lacZ* and *neo* cassettes to generate the floxed allele, *Rasa3*^*fl*^. *Rasa3*^*fl*/+^ mice were bred with *CMV-Cre* mice to generate mice carrying a germline null allele (*Rasa3*^+/−^). The *CMV-Cre* transgene was subsequently bred out by backcrossing to B6J mice. Crosses between *Rasa3*^*fl*/+^ and *Rasa3*^+/−^ mice generated *Rasa3*^*fl*/−^ mice. *Rasa3*^*fl*/−^ and *Rasa3*^*fl/fl*^ mice were bred to Cre-expressing transgenic mice to generate cell type and tissue specific knockouts.

### Polyinosinic-polycytidylic (Poly(I:C)) treatment

Floxed *Rasa3* exon three was excised in adult (8-12 weeks of age) hematopoietic cells by inducing *Mx1-Cre* recombinase activity with Poly(I:C) (GE Healthcare). Mice carrying floxed *Rasa3* allele(s) with and without (controls) *Mx1-Cre* were injected intraperitoneally with 300 μg of Poly(I:C) every other day for a total of five doses(42).

### Flow cytometric analysis of HSPC and terminal erythropoiesis

Dissociated whole spleens and bone marrow cells were passed through 100 μM Nitex nylon mesh (Genesee Scientific Products) with phosphate-buffered saline containing 0.5% bovine serum albumin and 2 mM EDTA (PBS/BSA/EDTA). Cell suspensions were filtered using CellTrics 50 μM disposable filters (Sysmex America), centrifuged at 300 x g for 10 minutes at 4 °C, and resuspended in 3-4 mL of PBS/BSA/EDTA. Quantitative flow cytometric analysis of hematopoietic stem and progenitor cells (HSPCs) was performed as described [43] following staining with conjugated antibodies. Stem and myeloid (SMP) progenitor cells (Lin^−^Sca^−^Kit^+^CD34^+^CD16/32^lo^) cells from mice on the BALB genetic background were sorted as described [35]. For analysis of terminally differentiated erythroid precursors [26], single-cell suspensions prepared as above were depleted of CD45 positive cells using mouse CD45 microbeads (Miltenyi Biotec) prior to staining. Antibodies used in flow cytometric analyses are given in Table S11. Stained cells were analyzed using LSR II Analyzer or FACSAria Sorter II (BD Biosciences) flow cytometers and FlowJo v. 9.9.3 software.

### Flow cytometric isolation of CD45+, Ter119+, and Ter119− cells from bone marrow and spleen

Single cell suspensions of whole spleen or bone marrow were prepared as above. CD45 positive cells were separated using mouse CD45 microbeads (Miltenyi Biotec). Ter119 positive cells were then isolated from the CD45 negative fraction using mouse Ter119 microbeads (Miltenyi Biotec) and magnetically separated using LS columns according to the manufacturer’s protocol. Ter119 negative cells were obtained as the flow-through fraction. Cell counts were performed using a Countess II Automated Cell Counter (Life Technologies) and/or Attune NXT Acoustic Focusing Cytometer (Thermo Fisher). Fractionated cell populations were used in western blotting (described below) or standard PCR reactions to genotype DNA for the presence of wildtype, floxed and/or excised alleles (S2 Table).

### In vitro colony-forming assays

Identification of BFU-e and CFU-e formation from whole bone marrow and spleen single cell suspensions was performed as described [44].

### Quantitative real-time RT-PCR (qRT-PCR) of HSPCs

Hematopoietic stem cells and progenitors including an enriched population of mixed erythroid BFU-E and CFU-E were isolated by flow cytometric cell sorting as described [19]. Total RNA was prepared using the PureLink RNA Mini Kit (Invitrogen). cDNA was synthesized using the High-Capacity cDNA Reverse Transcription Kit (Applied Biosystems) and random decamer primers (Invitrogen). qRT‐PCR reactions (25 μL total volume) consisted of 10 μL of cDNA, 1.25ul dH2O, 12.5 μL of TaqMan Gene Expression Master Mix and 1.25 μL of primers and probes (TaqMan® gene expression assay, assay ID: *Rasa3*, MM00436272.M1; Applied Biosystems). Samples were prepared in triplicate. qRT-PCR was performed using a ViiA7 Real-Time PCR System (Applied Biosystems) (40 cycles at 95 °C for 15 sec and 60 °C for 1 min). The relative quantification of each gene was determined by the 2^−ΔΔCT^ method using control CT values and *Gapdh* (TaqMan® gene expression assay ID: Mn99999915_g1) expression for normalization.

### Peripheral blood analysis

Whole blood (250 μL) was collected via the retro-orbital sinus using dipotassium ethylenediaminetetraacetic acid (K_2_EDTA)-coated capillary tubes (Drummond Scientific) into BD Microtainer® K_2_EDTA coated tubes (Becton, Dickinson and Company). Complete blood counts (CBCs) were obtained using an automated hematology analyzer equipped with species specific software (Advia 120 Multispecies Hematology Analyzer, Siemens Healthineers). Peripheral blood smears were stained with modified Wright-Giemsa (Sigma-Aldrich).

For cytokine measurements in bone marrow fluid, samples were prepared as described [45] with minor modifications. Briefly, bone marrow was flushed from femurs (0.5 cm) with 500 μL PBS using a 27 gauge syringe, resuspended, and centrifuged (600g, 5 min) to remove cells. Total protein content in bone marrow fluid was quantified by the method of Bradford [46] (Pierce™ kit, ThermoFisher Scientific). Bone marrow fluid samples and serum samples were analyzed by the University of Maryland Cytokine Core Laboratory Elisa Services (http://cytokines.com/).

### Phlebotomy

Adult *Mx1-Cre* negative control mice were rendered anemic by daily collection of ~350 μL whole blood via the retro-orbital sinus followed by intraperitoneal injections of sterile normal saline (37°C) for four days. Complete blood counts were obtained on day five.

### Sodium dodecyl sulfate–polyacrylamide gel electrophoresis (SDS-PAGE), western blotting and active RAS pull-down assays

Preparation of hemoglobin-depleted peripheral red blood cell membrane ghosts and whole cell lysates, SDS-PAGE and western blotting were performed as previously described [12]. Membrane ghost preparations were performed using packed red cells after removal of the buffy coat and thus are composed of both mature red cells and reticulocytes. Purified mature red cell and reticulocyte fractions for western blotting and RAS-GTP pull down assays were obtained as described [12]. Briefly, reticulocytes were separated from mature red cells by positive selection using antirat microbeads (Myltenyi) according to manufacturer instructions following staining with rat antimouse CD71 (Abcam).

The c-terminal peptide RASA3 primary antibody was generated as previously described [12]. GTP bound RAS was detected using the Active RAS Pull-Down and Detection Kit (Thermo Scientific) according to the manufacturer’s directions.

### Histology

Tissues were fixed in Bouin’s fixative, embedded in paraffin, sectioned at 3 μM and stained with hematoxylin and eosin for routine pathological analysis.

### RNAseq

Total RNA was isolated using the PureLink RNA Mini Kit (Invitrogen) and libraries prepared using the KAPA mRNA HyperPrep Kit (KAPA Biosystems) according to the manufacturer’s instructions, as previously described [20]. Libraries were checked for quality and concentration using the D5000 ScreenTape assay (Agilent Technologies) and quantitative PCR (KAPA Biosystems), according to the manufacturer’s instructions. Single-end sequencing was performed on the HiSeq2500 System using TruSeq SBS Kit v4 reagents (Illumina). Sequence reads were aligned and quantified with the Jackson Laboratory’s Computer Science group’s Civet Single-end RNAseq Analysis Pipeline (https://github.com/TheJacksonLaboratory/civet/). Briefly, sequences were preprocessed with an in-house python script that filtered sequences with less than seventy percent of the bases passing a quality threshold of 30, trimmed end bases with quality score less than 30, and removed sequences where the remaining length was less than seventy percent of the starting length. The resulting reads were mapped to mouse transcriptome (ENSEMBL version GRCm38.84) using bowtie *v2.2.0* (default parameters) and subsequent expression estimates were obtained using *rsem-calculate-expression v1.2.19* (default parameters). Upper quartile normalization of the resulting gene-level counts was accomplished with an in-house script. Count matrix normalization (rlog transformation) and differential expression analysis was performed with DESeq2 [47]. Differential expression analysis was performed on unnormalized counts assigned to each gene rather than FPKM (Fragments Per Kilobase of gene per Million reads), as recommended for DESeq2. Differentially expressed genes were extracted from DESeq2 output files using threshold values of ≥2 absolute fold change and ≤0.05 adjusted p-value.

To examine global patterns, further analysis was performed on the rlog regularized matrix. The regularized matrix was Z-transformed by the function *Z* = (*exp* − *avg*)/*stdev*, where *avg* and *stdev* are the average and sample standard deviation of the regularized expression level of each gene across all samples. To focus the analysis on genes with appreciable expression and/or variation, the matrix was reduced from all 38,925 annotated genes to 12,029 with *avg* ≥ 6.0 or *stdev* ≥ 2.0. (Similar reductions were made on tissue or cell-type specific Z-transformed matrixes for more specific Principal Components Analysis).

Principal Components Analysis was performed using default parameters with JMP v11.2.1 (SAS Institute) on the transpose of the reduced Z-transformed expression matrixes. Gene ontology (GO) term searches were performed with DAVID v6.8 (default settings) [36]. In some instances, redundant terms were removed in order to highlight additional processes.

### Image acquisition and assembly

Whole embryo images were acquired using a Nikon SMZ1500 stereomicroscope. Images of peripheral blood and tissue sections were acquired using a Nikon Eclipse E600 microscope equipped with a Nikon DS-Fi3 microscope camera and NIS-Elements Documentation software (Nikon Instruments). Embryos were examined with a 0.5X objective; smears with a 100X/1.4 oil-immersion objective; and tissue sections with a 20X/0.60 or 40X/0.75 objectives. Agarose gel thermal images and western blot films were scanned using a HP Scanjet G4010 scanner and HP Easy Scan software v.1.8 (Hewlett Packard). Final images were assembled using Microsoft PowerPoint for Mac v.16 (Microsoft).

### Image acquisition and assembly

Whole embryo images were acquired using a Nikon SMZ1500 stereomicroscope. Images of peripheral blood and tissue sections were acquired using a Nikon Eclipse E600 microscope equipped with a Nikon DS-Fi3 microscope camera and NIS-Elements Documentation software (Nikon Instruments). Embryos were examined with a 0.5X objective; smears with a 100X/1.4 oil-immersion objective; and tissue sections with a 20X/0.60 or 40X/0.75 objectives. Agarose gel thermal images and western blot films were scanned using a HP Scanjet G4010 scanner and HP Easy Scan software v.1.8 (Hewlett Packard). Final images were assembled using Microsoft PowerPoint for Mac v.16 (Microsoft).

### Statistical analysis

Significant differences were identified using Tukey honestly significant differences (HSD) or two-tailed Student *t* test using JMP v. 11 software (SAS Institute).

### Data Availability

RNAseq data were submitted to Gene Expression Omnibus (https://www.ncbi.nlm.nih.gov/geo/) with accession number GSE148821.

## Supporting Information

**S1 Fig. Breeding strategy to generate *Rasa3* alleles.** Schematic (adapted from Skarnes *et al.*)(41) showing strategy used to generate various *Rasa3* alleles. Targeted mutation (tm) designations used by convention by the KOMP (e.g., tm1a, tm1b, tm1c, and tm1d for the conditional ready targeted, germline null, floxed, and conditional null alleles, respectively) are indicated in blue. cKO, conditional knockout.

**S2 Fig. Germline RASA3 null mice are embryonic lethal**. Mice carrying the *Rasa3*^tm1a(KOMP)Wtsi)^ allele were produced on the inbred C57BL/6NJ (B6NJ) background by the KOMP at The Jackson Laboratory(48). Germline homozygous null mice generated by breeding with *Sox2-Cre* expressing transgenic mice (B6N.Cg-*Edil3^Tg(Sox2-cre)1Amc^*/J) die at E12.5-E13.5 with a phenotype of severe hemorrhage, overall pallor and a small, pale fetal liver (right, arrow). Additional data and images are available at the International Mouse Phenotyping Consortium (IMPC) website (www.mousephenotype.org).

**S3 Fig. Deletion of floxed alleles by *Epor-Cre* does not result in anemia. (A)** Genotyping in Cd45− Ter119+ spleen red cell precursors confirms excision of *Rasa3* floxed alleles in the presence of *Epor*-*Cre*. **(B)** Spleen weights as percent body weight (% BW). n = 3 per group. ns, not significantly different (p = 0.78). **(C)** Peripheral blood morphology is normal. A *scat* homozygote is shown for comparison. Bar, 10 μM. **(D)** Western blot of red cell membranes showing persistence of RASA3 protein in mutant mice. NS, non-specific band.

**S4 Fig. Frequency of hematopoietic stem and progenitors. (A)** Bone Marrow. **(B)** Spleen. LT-HSC, long term hematopoietic stem cell; ST-HSC, short term hematopoietic stem cell; LMPP, lymphoid-primed multipotent progenitor; CLP, common lymphoid progenitor; CMP, common myeloid progenitor; GMP, granulocyte monocyte progenitor; MEP, Myeloid-erythroid progenitor. n = 8 per group. * p < = 05.

**S5 Fig. Spleen histology. (A)** *Mx1-Cre*; *Rasa3* control and **(B)** mutant spleen. Effacement of the normal splenic nodular architecture and increased megakaryocytes are evident in the mutant spleen within 2 weeks of pIpC treatment. Original magnification 100x.

**S6 Fig. Spleen histology. (A)** B6J-+/+ **(B)** B6J-*hlb381/hlb381.* Original magnification 100x.

**S7 Fig. Analysis of global expression in bone marrow.** Hierarchal clustering **(A)** and principle component analysis **(B)** of expression differences in bone marrow.

**S8 Fig. Functional gene ontology (GO) terms for differentially expressed genes**. Data for all mutants (combined cr and pr) vs. WT in **(A)** spleen and (**B)** bone marrow. David BP_FAT function.

**S1 Text. References accompanying supplemental figures**

**S1 Table. Mouse strains**

**S2 Table. Genotyping primers and product sizes**

**S3 Table. Complete blood counts in *Epor-Cre; Rasa3* adult mice 6-8 weeks of age**

**S4 Table. Complete blood counts in *scat* mice 3-5 weeks of age**

**S5 Table. Complete blood counts in phlebotomized (PHB) control vs. mutant mice**

**S6 Table. Complete blood counts in B6NJ *Mx1-Cre; Rasa3* mice**

**S7 Table. Complete blood counts in C57BL/6J-*hlb381* mice 3-4 weeks of age**

**S8 Table. Complete blood counts in C57BL/6J-*hlb381* mice ≥ 6 weeks of age**

**S9 Table. RNAseq biological replicates**

**S10 Table. Complete blood counts in *scat* mice used for RNAseq**

**S11 Table. Flow cytometry antibodies**

**S12-24 Tables. Differential gene expression comparisons**

## Acknowledgements

This work was supported by National Institutes of Health Public Health Service grants HL134043 (L.L.P.) and P30CA034196 (The Jackson Laboratory NCI Cancer Center). We gratefully acknowledge the contribution the Genome Technologies, Flow Cytometry, Computational Sciences, and Histopathology Services at The Jackson Laboratory for expert assistance with the work described in this publication.

This work is dedicated to the memory of Dr. Barry Paw, an outstanding scientist and wonderful colleague and friend who left us too soon.

## Author contributions

**Conceptualization:** Luanne L. Peters

**Data Curation:** Raymond F. Robledo, Joel H. Graber, Luanne L. Peters,

**Formal Analysis:** Joel H. Graber, Nathaniel J. Maki, Lionel Blanc, Luanne L. Peters

**Funding Acquisition:** Luanne L. Peters, Lionel Blanc

**Investigation:** Raymond F. Robledo, Steven L. Ciciotte, Yue Zhao, Amy J. Lambert, Babette Gwynn

**Methodology:** Raymond F. Robledo, Luanne L. Peters

**Project Administration:** Raymond F. Robledo, Steven L. Ciciotte, Luanne L. Peters

**Supervision:** Raymond F. Robledo, Steven L. Ciciotte, Luanne L. Peters

**Validation:** Raymond F. Robledo, Steven L. Ciciotte, Luanne L. Peters

**Visualization:** Raymond F. Robledo, Luanne L. Peters

**Writing – Original Draft:** Luanne L. Peters

**Writing – Review and editing**: Lionel Blanc, Joel H. Graber, Raymond F. Robledo, Babette Gwynn, Luanne L. Peters

